# An inexpensive semi-automated sample processing pipeline for cell-free RNA

**DOI:** 10.1101/2020.08.18.256545

**Authors:** Mira N. Moufarrej, Stephen R. Quake

**Affiliations:** Department of Bioengineering, Stanford University, Stanford, CA USA 94305; Chan Zuckerberg Biohub, Stanford, CA USA 94305

## Abstract

Despite advances in automated liquid handling and microfluidics, preparing samples for RNA sequencing at scale generally requires expensive equipment, which is beyond the reach of many academic labs. Manual sample preparation remains a slow, expensive, and error-prone process. Here, we describe a low-cost, semi-automated pipeline to extract cell-free RNA (cfRNA) that like many RNA isolation protocols, can be decomposed into three subparts – RNA extraction, DNA digestion, and RNA cleaning and concentration. RT-qPCR data using a synthetic spike-in confirms comparable RNA quality as compared to manual sample processing, the gold-standard used in our prior work. The semi-automated pipeline also shows striking improvement in sample throughput (+12x), time spent (−11x), cost (−3x), and biohazardous waste produced (−4x) as compared to its manual counterpart. In total, this protocol enables cfRNA extraction from 96 samples simultaneously in 4.5 hours; in practice, this dramatically improves time to results as demonstrated in our recent work [1] where it was used to process 404 samples in 27 hours. Importantly, any lab already has most of the parts required (manual pipette, corresponding tips and kits) to build a semi-automated sample processing pipeline of their own and would only need to purchase or 3D-print a few extra parts ($5.5K total). This pipeline is also generalizable for many nucleic acid extraction applications, thereby increasing the scale of studies, which can be performed in small research labs.

## Introduction

Liquid biopsies offer an unprecedented opportunity to study human disease on a molecular scale and sample from the entire human body at once [2, 3, 4, 5, 6, 7, 8]. Applications of this technique span vast spaces like oncology [9, 10, 11], transplant care [12, 13], and prenatal diagnostics [14, 15, 1]. Numerous groups and companies have now demonstrated the utility of “omic”-based blood analysis in proof-of-concept work [6, 10, 13, 16, 17, 18]. However, scaling such discovery research remains obstructed by the lack of robust sample processing, which is paramount to yield trustworthy results – especially for human samples that may come from sources with disparate collection and storage guidelines.

Most discovery-focused work has four main stages – sample collection, sample processing, data generation, and data analysis. Sample collection, data generation, and data analysis have progressed tremendously due to the advent of mature biobanks, massively parallel sequencing (e.g., NovaSeq), and cloud-based computing (e.g., Amazon Web Services, Google Cloud) respectively. Sample processing however has remained manual despite increasing sample numbers. As a result, this laborious step constrains the number of samples processed at once and can lead to batch effects.

Although the biotechnology industry has long moved to robotic liquid handling solutions [19], many labs have not. Most automated solutions for lab work available today focus on production quality pipelines and do not accommodate research-based lab work in at least one of these ways – (1) No high-volume pipetting (e.g., Bravo), (2) No easy user-level customization (e.g., Hamilton, Eppendorf), (3) High initial expense ($20-100K) and custom consumables (e.g., Bravo, Hamilton, Eppendorf).

In this protocol, we describe the design and validation of a semi-automated pipeline to extract cell-free RNA (cfRNA) by outfitting an existing inexpensive, open-source robotic platform, the Opentrons 1.0 (OT1). Specifically, we detail how to setup and calibrate the robotic system, validate the platform end-to-end using an exogenous RNA control, and finally use the system to process clinical samples in high-throughput with suitable quality for downstream analysis [1]. We also show that RT-qPCR data highlight comparable results to the gold-standard used in prior work - manual sample processing [15]. Overall, this protocol provides striking improvement in sample throughput (+12x), time spent (−11x), cost (−3x), and biohazardous waste produced (−4x) as compared to its manual counterpart, and enables cfRNA extraction from 96 samples simultaneously in 4.5 hours. Importantly, any lab already has most of the parts required (manual pipette, corresponding tips and kits) to build a semi-automated sample processing pipeline of their own, and would only need to purchase or 3D-print a few extra parts ($5.5K total). Such robust and scalable solutions will allow us to test hypotheses faster and spend less time on repetitive but important tasks like sample processing.

## Experimental design

### Robotic system overview

Like many RNA isolation protocols, processing cfRNA samples can be decomposed into three subparts – RNA extraction, DNA digestion, and RNA cleaning and concentration (Fig. 1A). Our process requires a single robotic system than can easily fit within a chemical hood – necessary to process bodily fluids – and can be configured to use any individual eight-channel pipette. Nearly all steps are performed on the robot with minimal user input. At the system’s prompting, the user fills reagent boats, transfers plates for centrifugation, or swaps out tip boxes. Sample addition is performed by hand to easily accommodate various sample storage strategies (e.g. 1.5 mL microcentrifuge tubes, matrix tubes). Similarly, DNAse master mix and eluant addition are also performed by hand to maintain the system’s simplicity since these steps stand out as the only small volume (*<*15 *µ*L) pipetting in the process. By using standard lab equipment and interfacing with the user at key steps, our semi-automated pipeline streamlines sample processing while maintaining flexibility key to discovery stage research.

**Figure 1:**
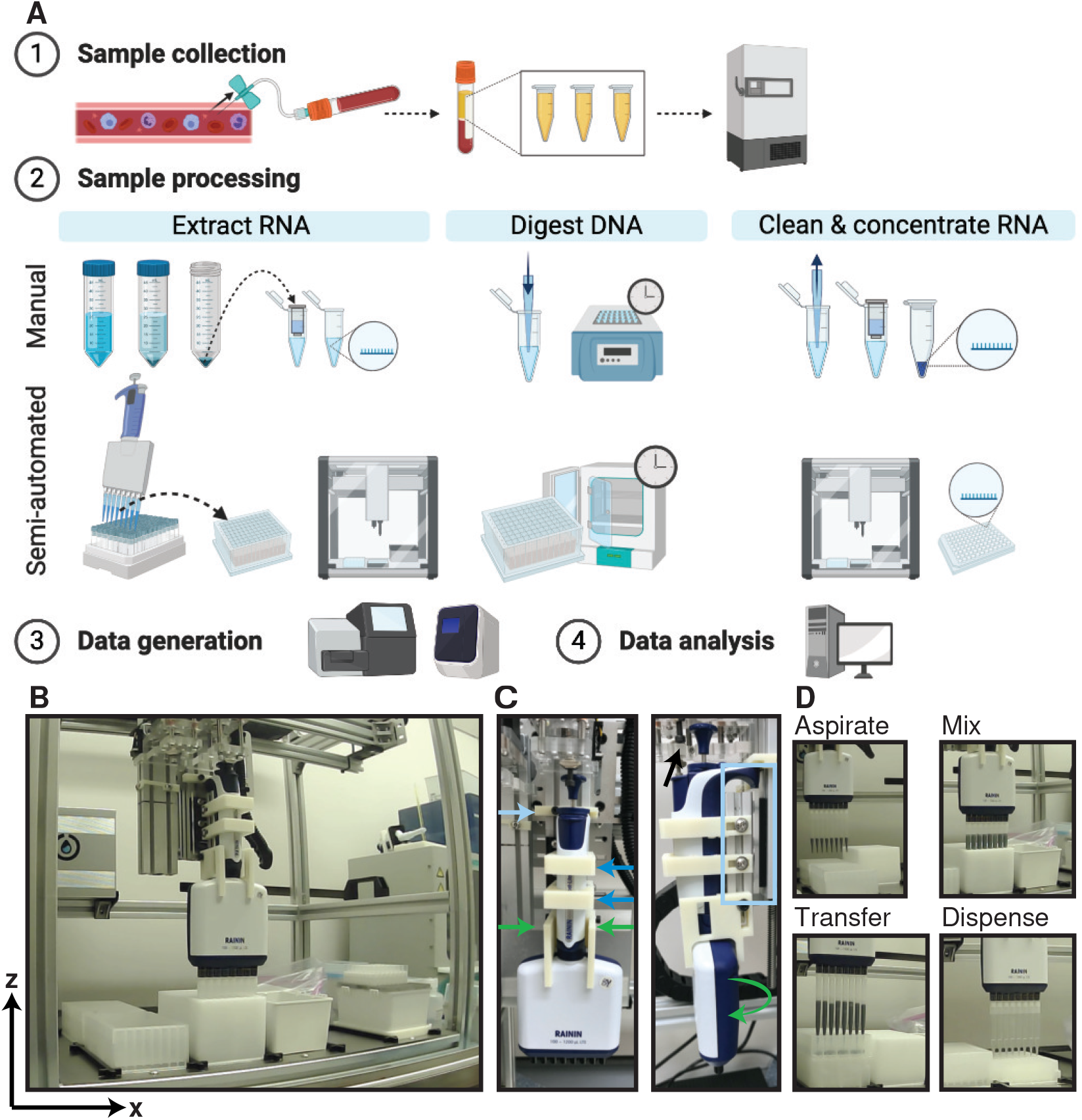
Overview of semi-automated RNA isolation system. (A) Graphical overview of sample collection, processing, data generation and analysis of cfRNA data. Step 2 compares manual and semi-automated sample processing. (B) Representative photograph of full system. (C) Close-up of mounted eight-channel high volume pipette. Dark blue, green, and light blue indicators highlight 3D printed fixtures to prevent motion in the x, y, and z directions respectively. Black arrow indicates moving fixture used to eject tips. (D) Representative freeze-frame photographs of various tasks performed on the robotic system.

### Robotic system setup

The OT1 system is composed of motors that drive motion in the x, y, and z directions across 10 deck positions where the user can place reagent boats, plates, tip boxes, and trash (Fig. 1B). To use it, the user must mount their pipette of choice so that it is rigidly fixed in all directions and only moves with the motors themselves. We achieved this by designing and 3D-printing several parts to work with our pipette of choice. To fix motion in the x-direction, we mounted two U-shaped clasps (Fig. 1C, dark blue arrows). These clasps are then fastened to a hard back (Fig. 1C, light blue box). This back board coupled with a piece mounted on top of the pipette (Fig. 1C, light blue arrow) prevents unexpected z-axis motion during tip retrieval and disposal. Finally, two more L-shaped holds (Fig. 1C, green arrows) prevent y-axis motion in the pipette body, which can otherwise rotate freely. Together, these 3D-printed fixtures allow the system to remain properly calibrated across multiple runs while mounting and disposing of tips to aspirate, mix, transfer, and dispense reagents (Fig. 1D, 2, Full protocol at https://youtu.be/g6RsSaNvSNA). Additionally, we designed and printed a holder for pipette tip boxes to fit within a standard deck footprint. For more details regarding pipette installation, see https://support.opentrons.com/en/articles/689945ot-one-installing-pipettes.

**Figure 2:**
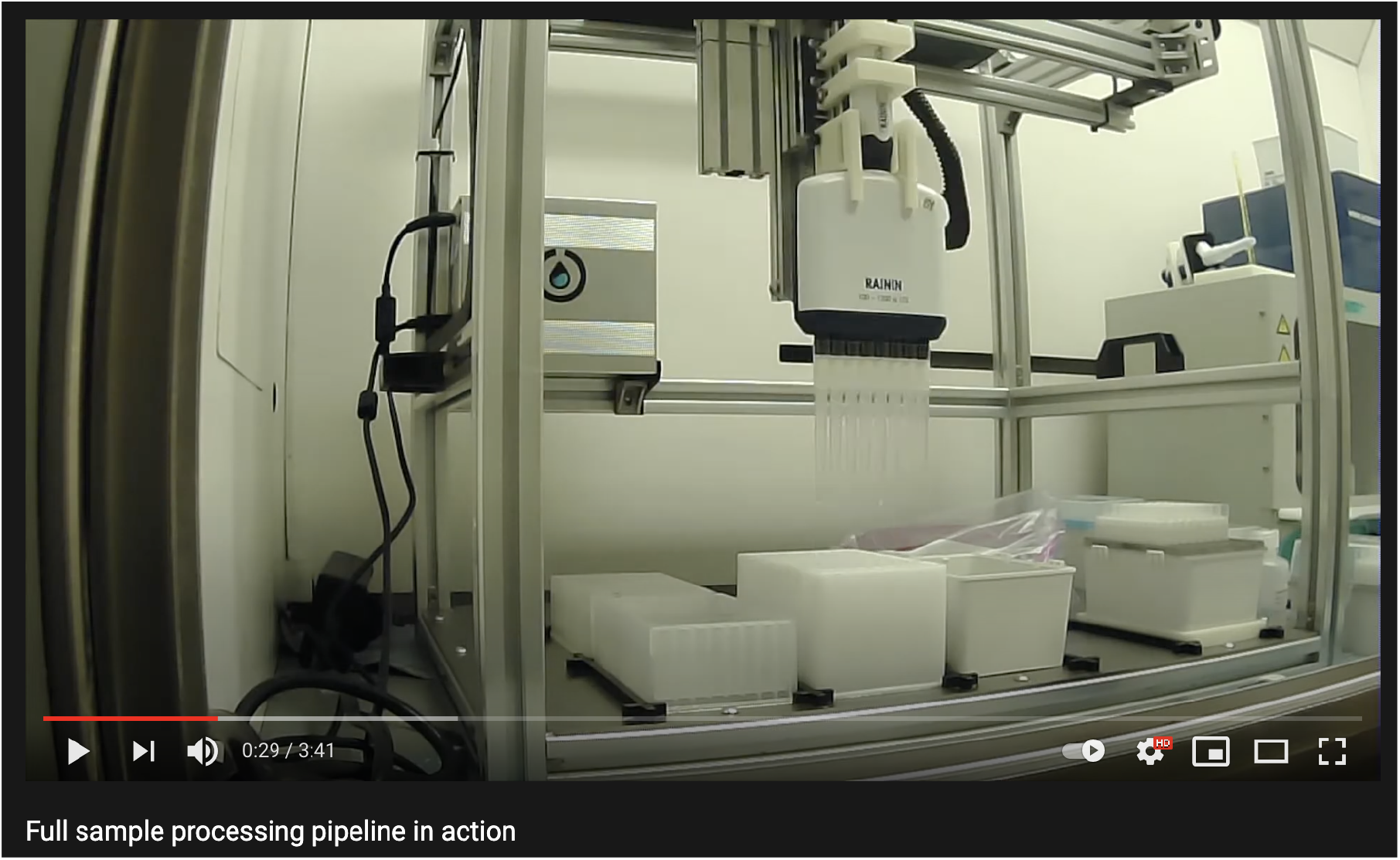
Full-protocol at 20x speed. Slides in white indicate user steps like centrifugation. Screenshot included above, See https://youtu.be/g6RsSaNvSNA for full video.

### Robotic system calibration

Once the pipette has been fixed onto the system, it is necessary to calibrate the OT1, and register the x,y, and z position for each consumable in each deck position it may occupy. To this end, we have provided a series of calibration scripts and detailed instructions at https://github.com/miramou/cfRNA_pipeline_automation/tree/master/Opentrons/scripts/calibration that can be used with the OT1 software to calibrate the system. Special care should be taken when calibrating the pipette tip boxes as even small miscalibrations may result in a failure to pick up tips or faulty tip placement with negative downstream consequences like mispipetting. More details can be found at https://support.opentrons.com/en/articles/689977-otone-calibrating-the-deck.

### Protocol overview and validation

The reagent volumes required to isolate cfRNA depend on initial sample volume (typically 1 mL for our process). To extract cfRNA from 1 mL of plasma requires 5 mL of added reagents yielding a maximum total volume of 6 mL. Since this is far beyond what a typical 96-well deep well plate can accommodate (about 1-2 mL), we opted to use 48 deep-well plates to maintain system scalability (Fig. 1A). Using a 48 deep-well plate not only interfaces well with the robotic system, but it also allows us to move away from the classic 50 mL conical tubes used for this step, since these can be difficult to process quickly in an automated fashion (Fig. 1A). To process 96 samples using 48-well plates requires that the samples are split into two batches briefly and then recombined into 1 plate for subsequent steps (Fig. 3A). Specifically, we thaw and extract cfRNA from 48 samples (Fig. 3A left side) and then refrigerate the isolated cfRNA while thawing and processing cfRNA from another 48 samples (Fig. 3A right side).

**Figure 3:**
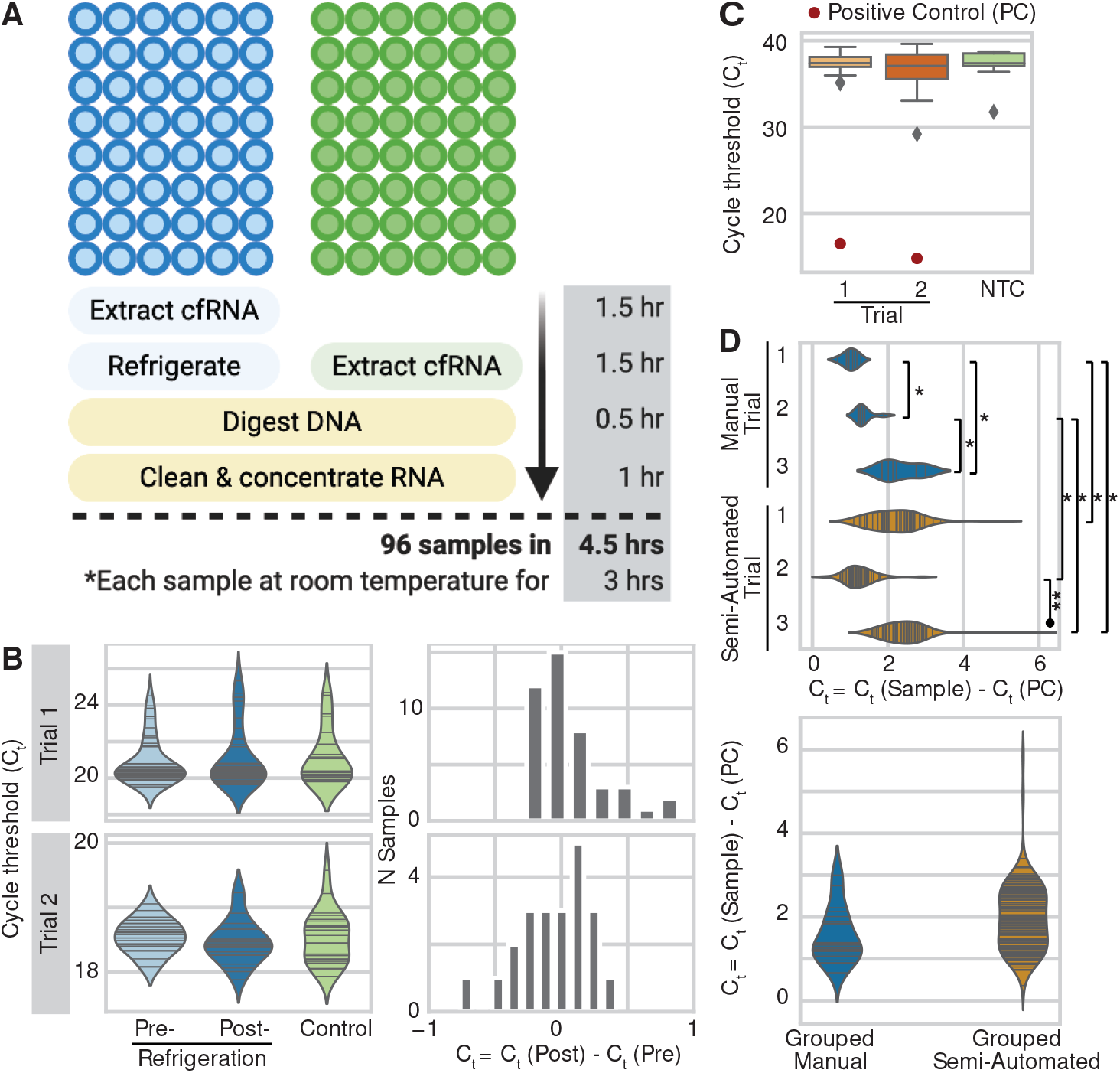
Semi-automated RNA isolation system validation using RNA control. (A) Graphical overview of semi-automated sample processing. (B) Comparison of RT-qPCR cycle thresholds (Ct) for paired samples pre- and post-refrigeration (light and dark blue) as compared to unrefrigerated samples (green) for two independent trials (rows). Histograms (right column) further highlight the distribution of difference in Ct between paired samples post- and pre-refrigeration. (C) Boxplot to identify any cross-contamination during processing. Yellow and orange boxplots show the distribution of Ct for processed water samples from two independent trials as compared to those for known non-template controls (NTC) in green. Red points indicate positive controls. (D) Comparison of manual and semi-automated protocols with three trials per method. Top and bottom violin plot compare all trials in either an independent fashion or grouped together respectively. Symbols * and ** indicate statistically significant differences between the two indicated groups at thresholds less than of 0.05 and 10^*−*10^ respectively. 36

We investigated whether this refrigeration step had adverse effects on cfRNA concentration by spiking in a known concentration of a single RNA oligonucleotide at the start of extraction. We then measured its concentration using RT-qPCR pre- and post- refrigeration as compared to that for control samples processed without any refrigeration (Fig. 3B left). Across 2 independent trials, we find that refrigeration of isolated cfRNA for 1.5 hours has no adverse effects on final yield (Bonferroni adjusted p-value = 0.43 and 1.0 per trial, Wilcoxon signed-rank test) (Fig. 3B right). Further, we find no loss in yield for samples post-refrigeration as compared to control unrefrigerated samples (Bonferroni adjusted p-value = 1.0 and 1.0 per trial, one-sided Mann-Whitney rank test). We also separately confirmed that the custom RNA oligonucleotide and its corresponding TaqMan probes work as expected (Fig. 4).

**Figure 4:**
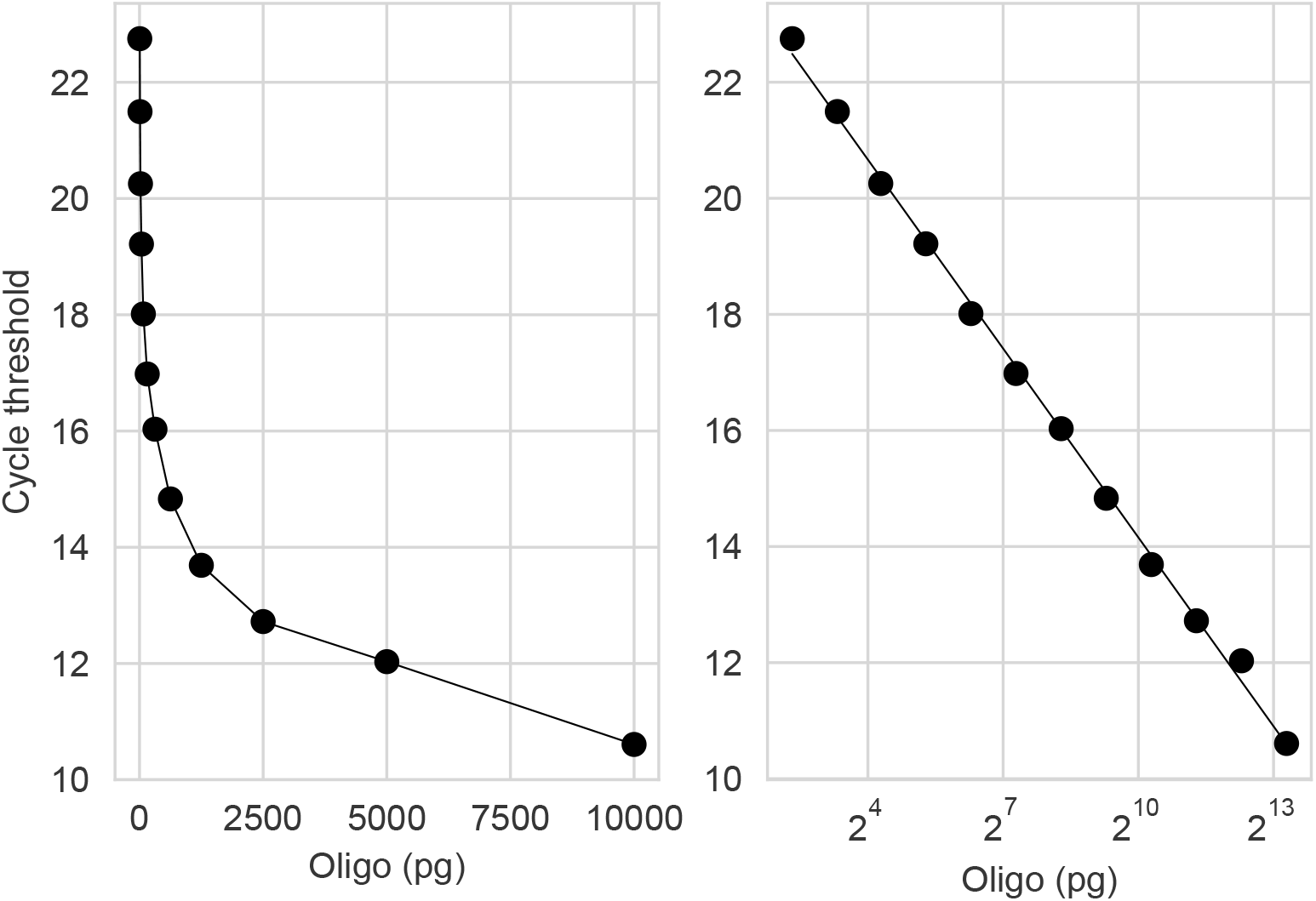
Validation of custom RNA control, ERCC54, and its corresponding probes across a range of oligonucleotide input values. Left and right plots show linear and log x-axis values respectively.

Subsequently once cfRNA from all 96 samples have been extracted, all samples are treated with DNAse to remove any lingering DNA, cleaned, and concentrated to a final volume of 12 *µ*L (Fig. 3A). In total, this process takes 4.5 hours with each sample at room temperature for 3 hours at a given time (Fig. 3A). Finally, we quantified the likelihood of cross-contamination between samples by spiking in RNA only into every other well so that each 48-well plate then contained 24 sample wells and 24 blank wells.

Across 2 independent trials with 48 blank wells each (24 per 48-well plate), we find no evidence of cross-contamination (Fig. 3C). Specifically, we find that blank samples do not yield a distribution of RT-qPCR cycle thresholds (C_*t*_) significantly lower than that of true non-template controls (NTC, n = 17) (Bonferroni adjusted p-value = 0.16 and 0.10 per trial, one-sided Mann-Whitney rank test). Positive controls (PC) are included for reference (Fig. 3C).

When setting up the system and periodically when processing samples regularly, we recommend that the user confirms the absence of cross contamination prior to processing precious clinical samples.

### RT-qPCR validation

To compare the semi-automated pipeline to the gold standard of traditional manual pipetting, we compared the concentration of RNA rescued for a known control relative to the concentration of RNA expected with total rescue (∆ C_*t*_). Over 3 independent trials per protocol, we find that the manual and semi-automated methods perform comparably (Fig. 3D).

Globally, when we combined ∆C_*t*_ values from all 3 trials per protocol, we found that extracting RNA using the semi-automated method (n = 144, 48 per trial) did not significantly decrease RNA rescue when compared to the manual protocol (n = 24, 8 per trial) (Bonferroni adjusted p-value = 0.11, one-sided Mann-Whitney rank test). However, when comparing individual trials in a pairwise fashion, we observed some significant shifts to more positive ∆C_*t*_ values specifically between all 3 manual (M) trial pairs, 1 semiautomated (SA) trial pair, and 4 of 9 manual and semi-automated trial pairs (Bonferroni adjusted p-value = 0.031, 0.0075, 0.016, 1.83 × 10^*−*14^, 0.0001, 5.86 × 10^*−*5^, 0.002, 0.00013 for pairs M1:M2, M1:M3, M2:M3, SA2:SA3, M1:SA1, M1:SA3, M2:SA1, M2:SA3 respectively, one-sided Mann-Whitney rank test). We note that because the manual trials (n=8) require smaller batch sizes than their semi-automated counterparts (n=48), pairwise comparison differences may be the result of sub-sampling rather than differences in protocol yield. The lack of a significant difference under global testing conditions supports this theory.

## Comparison to other methods

### Cost, time, and sustainability comparison

When compared to a manual protocol assuming a batch size of 8 samples (Fig. 1A), semi-automated parallel processing results in a 12-fold increase in sample batch size and nearly an 11-, 3-, and 4-fold reduction in time, cost, and biohazardous waste produced respectively (Table 1). We note that this RNA isolation protocol like many of its kind lends itself well to parallelization because incubation and centrifugation compose nearly half of total processing time. As sample number increases, such steps remain near constant in time. Further, reduced plastic usage, specifically by shifting to plate-based (vs individual tube) consumables and reduced tip usage, majorly contributed to decreased cost and biohazardous waste produced per sample processed.

**Table 1:** Comparison of manual and semi-automated RNA isolation protocols.

| Method | Batch size<br>(n) | Time per batch<br>(hours) | Cost per sample<br>(\$) | Tips used per<br>sample (n) |
| --- | --- | --- | --- | --- |
| Manual | 8 | 4.5 | 41.25 | 26 |
| Semi-automated | 96 | 4.5 | 15.24 | 7 |

### Advantages and limitations of the protocol

The key benefits of a semi-automated system including an increase in sample throughput by 12-fold and reduction in time spent, cost, and biohazardous waste produced by nearly 11-fold, 3-fold, and 4-fold respectively. Such quantitative benefits are coupled with several qualitative improvements distinct from what one can expect from a manual protocol. Specifically, a semi-automated protocol allows for consistent sample preparation quality across users, reduced personnel time, and reduced user concentration during the protocol. When contrasted with fully automated systems, we find that a semi-automated system is well-suited to the needs of a typical academic lab performing discovery research. Although a semi-automated system requires the user to remain nearby, its initial cost ($5.5K) remains significantly cheaper than its automated counterparts ($20-100K).

Importantly, the system’s low cost and easy user-level customization can be applied to automate any RNA isolation required for discovery research in an academic lab. Because the system is specifically built to leverage existing 96-well kits for RNA isolation, cleaning, and concentration, such a protocol can easily be transferred to isolate RNA from other sources like tissues, urine, or other bodily fluids with relatively few changes. The user would only have to program the new liquid handling steps or modify existing ones using a common high-level programming language (Python). This easy customization stands in stark contrast to the proprietary software or lower-level programming languages like C required to modify protocols on most automated systems.

The semi-automated protocol we describe is not without its limitations – namely that the OT1 system selected here does not permit for easy pipette changes restricting the user to a single multi-channel pipette’s volume range. In an academic lab setting, we do not see volume restrictions as a major issue since most RNA protocols require reagent volumes within the range of a single pipette. The user can perform any steps that require reagent volumes outside the range of the mounted pipette by repeatedly transferring a smaller volume within the upper limit for larger volumes or transferring by hand for smaller volumes. We find performing the three steps that require smaller, variable volumes by hand provides additional flexibility in the protocol. For instance, adding samples by hand easily accommodates various sample storage strategies like microtubes and matrix tubes. Additionally, adding the eluant by hand permits the user to easily modify the volume used for each RNA isolation to match their chosen downstream application.

We also found that setting up and calibrating manual pipettes to work with this platform required significant effort. These pain-points have since been addressed in the second generation of this robotic system, the OT2 ($6.5K including pipettes), which can be used in place of the OT1 system described here. Users who choose to use the OT2 may skip the section above titled “Robotic system setup” and instead follow the directions provided at the Opentron’s website regarding setup and subsequently proceed immediately to calibration. Additionally, the Python API to interface with the OT2 as compared to the OT1 may be slightly different and some function calls may have to be tweaked. The general liquid handling logic flow however will remain consistent, and we expect these changes to be minor. Although the OT1 since its use in this work has been phased out, all necessary hardware and software required to independently build the system have been made available on GitHub [20]. Further, both systems are open-source permitting users to easily and inexpensively tinker to exactly suit their needs.

Overall, we have shown that semi-automated sample processing can be performed affordably and quickly within individual academic labs. Shifting to semi-automated sample processing alleviates a significant bottleneck in discovery-focused work and allows for seamless transitions from sample collection to data generation and analysis. With this robust and scalable solution, we can test hypotheses faster and spend less time on repetitive but important tasks like sample processing.

### Applications

The current protocol is suitable for plasma samples (preferred, fresh or previously frozen at −80°C) or serum samples. An example bash script to launch the protocol is provided for 0.75 and 1 mL volume samples. These can be easily adapted to accommodate samples of different volumes following the instructions provided by Norgen, which manufactures the RNA extraction kit used here.

### General considerations

This protocol details how to extract cfRNA simultaneously from 96 samples using a robotic system. We provide resources to guide users to set up and calibrate a robotic system including a list of parts, specific scripts for calibration, and photographs and videos of a representative well-calibrated system. Manual pipetting can be substituted for robotic pipetting and may be preferred when learning or troubleshooting the protocol.

Our protocol is written for researchers with some familiarity with the command line and programming languages like Python and Bash. The user can also choose to run the protocol through Opentron’s provided software to interface with the robot, but should be aware that such a transition will require some modification of the existing codebase and require that the user understands the protocol enough to manually transition between sub-protocols, which would otherwise occur automatically when run using the command line.

Prior to processing samples, it is advised to map out the plate and randomize samples across it. If samples from a case/control study are processed, confirm that each half plate (48 samples) contains samples from both case and control subjects to ensure that any batch effect can be decoupled from the case effect.

### Considerations about the laboratory facilities

All the steps in our protocol should be done in a standard laboratory with a chemical hood. Prior to starting, care should be taken to ensure the workspace and all consumables placed within it are clean (e.g., using 70% EtOH) and RNase-free (e.g., using RNase Away).

## Materials

### Biological materials

- Fresh or frozen plasma samples
  · **CRITICAL:** Ensure that sample collection adheres to your local IRB or APLAC guidelines. Tubes used for plasma isolation can contain most common solutions with the exception of heparin, which inhibits reverse transcription downstream. More details about best practices can be found in [21] (see section titled “Experimental challenges when isolating circulating RNA”).

### Reagents

- Norgen Plasma/Serum Circulating and Exosomal RNA Purification 96-Well Kit (Slurry Format) (Norgen, cat. No. 29500)
- Baseline-ZERO DNAse (Lucigen, cat. no. DB0715K)
- Zymo RNA Clean and Concentrator-96 kit (Zymo Research, cat. no. R1080)
- RNA 6000 Pico Kit (Agilent, cat. no. 5067-1513)
- 2100 Bioanalyzer instrument (Agilent)
- 2-mercaptoethanol (Sigma-Aldrich, cat. no. M6250)
  · **CAUTION:** 2-mercaptoethanol is highly corrosive and toxic and poses a health and environmental hazard. Avoid skin contact or inhalation. Wear full personal protective equipment (PPE) when handling always within a chemical hood. Apply other safety measures if necessary, and adhere to local guidelines when disposing.
- 200-proof ethanol (Sigma Aldrich, cat. no. E7023)
- Nuclease free H_2_O (IDT, cat. no. 11-04-02-01)
- iTaq Universal Probes One-Step Kit (Biorad, cat. no. 1725141)
- SMARTer Stranded Total RNAseq Kit v2 - Pico Input Mammalian Components (Takara, Cat No 634419)
- SMARTer RNA Unique Dual Index Kit – 96U Set A (Takara, Cat No 634452)

### Automation compatible consumables

- 48-well deep well plates (Thomas Scientific, cat. no. 1149Q15)
- 12-column reagent reservoir (Thomas Scientific, cat. no. 1149Q70)
- 2-column reagent reservoir (Thomas Scientific, cat. no. 1149Q79)
- 1.2 mL pipette tips (Rainin, cat. no. 30389231)
- 1 mL extended length pipette tips (Rainin, cat. no. 30389223)
  · **CRITICAL:** Extended length tips are necessary to ensure the full sample volume is transferred at Step 38. If substituting for another pipette and its corresponding tips, one should ensure that the tips used are adequately long to reach the bottom of a 48-well deep well plate.
- 96-well PCR plate (Biorad, HSP9631)
- 2 mL deep well plates (Thomas Scientific, cat. no. 1149J85)

### Equipment

- 3D printer
- 3D printed parts (See Github)
- Opentrons 1.0 (OT1)
- 8-channel, 1.2 mL pipette, L8-P1200 (Rainin)
- 8-channel, 20 *µ*L pipette, L8-P20 (Rainin)
- Vortex
- Chemical hood
- Incubator
- Centrifuge
  · CRITICAL Centrifuge must achieve spin speeds of 3000-5000xg and include buckets for deep well plates.
- -80°C, −20°C freezer
- Refrigerator
- Real-Time PCR Detection System (Biorad, CFX96 or CFX384)
- Razor blades (Ted Pella, cat. no. 121-32)
- Ziploc gallon bags
- Aluminum foil seals (Sigma Adrich, cat. no. Z722634)
- Adhesive clear plate seals (Fisher Scientific, cat. no. AB-0558)

### Exogenous RNA control and probes

- Custom short oligonucleotide that matches 3’ end of External RNA Controls Consortium 54 (ERCC54)
  · rUrUrU rArGrA rArUrG rCrUrU rArArA rGrArU rGrGrC rArGrA rGrUrU rGrGrA rGrGrA rGrArG rArUrU rUrGrC rCrArA rUrCrA rCrArA rArCrC rArArA rUrCrA rGrUrU rGrArG rUrGrG rArGrG rGrCrA rArCrA rArCrA rGrArG rArGrU rUrGrC rUrArU rArGrC rGrArG rGrGrC rUrUrU rGrGrC rArArA rCrArA rCrCrC rArCrC
- TaqMan gene expression assay for ERCC54 (Thermo Fisher, cat. no. 4448490, Assay ID Ac03459999 a1)

### Software

- Opentrons app (v 2.5.2)
- Miniconda
- Git
- Python
- Bash
- Software for running the protocol is available at:
  · https://github.com/miramou/cfRNA_pipeline_automation/tree/master/Opentrons
- Software for processing raw data is available at:
  · https://github.com/miramou/cfRNA_pipeline
- Software to reproduce the figures in this manuscript is available at:
  · https://github.com/miramou/cfRNA_pipeline_automation/tree/master/Manuscript_Figs

### Reagent setup

- Filter plate balance (Norgen, Zymo) and 48-well plate balance: Ensure that you have balances for each of these plate types for centrifugation step along with appropriate water balances
- Exogenous RNA control: Reconstitute lyophilized RNA oligonucleotide per manufacturer instruction at a concentration of 1 *µg* per *µL*. On ice, aliquot stock solution into 2.5 *µL* single use aliquots. Store at −80°C indefinitely.
- Lysis buffer A (Norgen): add 1.2 mL 2-mercaptoethanol per 100 mL lysis buffer
- Wash buffers (Norgen and Zymo): Add 200-proof ethanol to wash buffer per manufacturer instructions
- Lysis buffer and slurry: Preheat in 60°C incubator, vortexing occasionally until all precipitate dissolves and slurry appears well combined.
  · **CRITICAL:** Residual precipitate may clog pipette tips during reagent transfer leading to inaccurate pipetting. Keep buffer and slurry heated until immediately before use, at which point, vortex once more.

### Equipment setup

- Clean and preheat incubator to 60°C
- Clean all work surfaces including OT1 robot deck, surrounding workspace in chemical hood, and labware with 70% EtOH and a RNase decontaminant to ensure a RNase free space.

### Software setup

- Follow the setup instructions provided under the heading “Getting started” at this link: https://github.com/miramou/cfRNA_pipeline_automation/tree/master/Opentrons. Specifically, the user will have to clone the repository and create a new miniconda environment using the provided requirements file.

### General setup

- If confirming RNA quality using the Bioanalyzer, select 11 wells at random across the plate spanning both columns and rows, to be run on 1 chip.

## Procedure

### Prepare samples

Timing: 10 minutes

1. Prior to thawing any samples, ensure that you have mapped out the plate (see General considerations). Of special note, if samples are associated with a case/control, confirm that each half plate (48 samples) contains samples from both case and control subjects to ensure that any batch effect can be decoupled from the case effect.
2. See option A or B for compatible sample types that we have tested. We strongly recommend starting with Option A as the user familiarizes themselves with the protocol. Once the user feels comfortable with the entire approach, Option B becomes the standard. We highlight that given the precious nature of clinical samples, we have found it important to thoroughly test the entire pipeline first.
  A. Exogenous RNA control
    i. On ice, thaw single-use aliquot of exogenous RNA control.
    ii. Once thawed, quickly spin down and dilute further to yield a solution at 5 ng/*µ*L by combining 1 *µ*L of stock with 199 *µ*L of nuclease-free water.
    iii. To mimic the expected sample volume of 1 mL, pipette 1 mL of nuclease-free water per well across 48 wells of a 96 deep-well plate on ice.
    iv. Add 1 *µ*L of diluted stock to each well of the stock plate and mix well. Seal and store at 4°C until needed for RNA extraction.
  B. Frozen plasma samples
    i. At room temperature, thaw the first 48 plasma samples. Do not thaw at 4°C to prevent the formation of cryoprecipitates that inhibit RNA extraction performance.
    ii. Organize samples to match planned plate layout so that they can be easily pipetted using a multi-channel pipette (see RNA extraction).
    iii. As samples thaw, proceed to prepatory steps for RNA extraction.

### RNA extraction

Timing: 3 hours (1.5 hours per 48-well plate)

Once the launch script starts and throughout the protocol, the system will also prompt the user about when and where to add reagents, swap tip boxes, and place consumables among other necessary cues. Note that the stated reagent volumes at steps 16 and 22 reflects the processing of 1 mL plasma samples. As noted below, the protocol can easily be adjusted to process other sample volumes. See Fig. 2, https://youtu.be/g6RsSaNvSNA for a full run-through of the protocol.

3 Turn on and connect the robotic system. Confirm that the computer recognizes the robotic system using the Opentrons app. Home the system.
4 Using the shell, deploy launch script titled `run_cfRNA_pipeline.sh` passing one argument that points to where the user would like log files saved.
  A. **CAUTION:** Sample launch script is programmed to process 1 mL samples. Other sample volumes require different reagent volumes. Refer to manufacturer instructions (Norgen Plasma/Serum Circulating and Exosomal RNA Purification 96-Well Kit) for these volumes, and modify global variables `LYSIS_VOL` and `ETOH_VOL` in launch script accordingly.
5 Seal one new 2 column reagent reservoir and one new 12 column reagent reservoir using a clear plate seal. The seal will be sequentially cut to prevent reagent cross-contamination.
6 Using a razor blade, cut the seal on the 12-column reservoir such that the boat labeled 1, which will be closest to the user once placed on the deck, is now exposed and part of boat 2 immediately adjacent to it is also open. Reagents will only be added to every other boat, specifically boats with odd numbers to further prevent reagent cross-contamination and prevent pipette tips from sticking to the seal when entering a boat.
7 Using a razor blade, cut the seal on the 2-column reservoir such that the boat labeled 1, which will be closest to the user once placed on the deck, is now exposed.
8 Place standard length 1 mL filter tip box (gray if using Rainin) at position E1 and completely remove lid.
  A. **CRITICAL:** If using a pipette box where the footprint does not match that of deck slot like Rainin boxes, 3D print and use a pipette box holder to fix its position. For Rainin, we provide a box holder design at this link: https://github.com/miramou/cfRNA_pipeline_automation/blob/master/Opentrons/custom_solidworks_parts/tip_box/OPENTRONS_RAININ_TIP_BOX_HOLDER_2.STL
9 Place a Ziploc bag with the sides rolled down halfway at position D2. This will act as trash where the robot will dispose of used tips. Ensure that the bag sits upright.
10 Place the 2-column reagent reservoir at position A1. Place the 12-column reagent reservoir at A2. Deck positions can be referenced in the Opentrons App.
11 Place a new 48-well deep well plate at position B2.
12 Vortex preheated slurry solution from the Norgen kit well, and add 10.5 mL to column 1 in the 12-column reservoir at A2, which should be closest to the user and fully unsealed.
  A. **CRITICAL:** Proceed quickly. In our experience, cooled slurry crystallizes and clogs the pipette tip upon reagent transfer.
13 Press enter to proceed with protocol. The robotic system will now pick up pipette tips and add 200 *µ*L slurry to each row of the 48-well plate and then dispose of used tips in the trash bag.
14 Move the 48-well plate to position B1 once the system prompts you.
15 Vortex preheated lysis buffer A from Norgen kit and add 110 mL to position 1 in the 2-column reservoir at A1, which closest to user and unsealed.
  A. **CRITICAL:** Proceed quickly. In our experience, cooled lysis buffer crystallizes and clogs the pipette tip upon reagent transfer.
16 Press enter to proceed with the protocol. The robotic system will now pick up pipette tips and add 1.8 mL lysis buffer to each row of the 48-well plate and then dispose of tips.
17 Using a new tip box, transfer samples by hand (see Option A or B above in “Prepare samples”) to the 48 well plate and mix each column by pipetting up and down at least 5 times. Set aside the tip box. You will use the same box for all 96 samples in two rounds.
  A. **CRITICAL:** Do not use the already opened tip box on the robot. The protocol is programmed to know how many rows of tips have been used and pick up where it left off. Additionally, if processing 96 samples, a single tip box can be entirely dedicated to sample transfer and set aside between batches of 48 samples.
18 Seal plate well and incubate at 60°C for 10 minutes.
19 Type ‘y’ and press enter to proceed with protocol.
20 Once there is approximately 1 minute left in incubation, remove the seal completely from the 2-column reservoir at position A1 and add 150 mL 200-proof EtOH to the 2nd column further from the front.
21 After the incubation is complete, remove the seal from the 48 well plate and place it at deck position B1.
22 Press enter to proceed with the protocol. The robotic system will now pick up pipette tips and add 3 mL 200-proof EtOH to each row of the 48-well plate and then dispose of used tips in the trash bag.
23 The system will now automatically proceed to mixing each row of samples by pipetting up and down at different heights and dispose of tips after each row to ensure that each 5 mL sample is well-mixed.
24 Once prompted, seal the 48-well plate well and centrifuge for 1 min at 2000 RPM to create a slurry pellet. Ensure centrifuge is appropriately balanced. Move the 48-well plate back to the chemical hood.
25 Place a hard-sided open vessel at position C1. This will serve as liquid trash as the robot decants supernatant from the 48-well plate. Typically, an empty tip box with the tip rack removed works well for this purpose.
  A. **CRITICAL:** When running through this protocol for the first time, ensure that pipette tips do not touch the liquid waste when decanting and that splatter is minimized. Recalibrate otherwise.
26 We will remove the seal sequentially from each row. Using a razor blade, cut the seal between columns 1 and 2 and remove the seal to reveal the 8 samples in column 1 alone. Place the plate at position B1 with column 1 closest to the user.
27 Press enter to proceed. The robot will now decant all supernatant from column 1 and then dispose of tips. If at any point, the trash bag containing tips becomes full, use the programmed pause between columns to also replace the trash bag with a new one.
  A. **CAUTION:** Dispose of biohazardous trash and 2-mercaptoethanol according to local environmental guidelines.
28 Repeat steps 26 and 27 for each column of the 48-well plate. After the 3rd column, the system will prompt you to replace the now empty tip box at E1 with a new tip box. Do so, taking care to remove the plastic lid completely from the tip box. As before, press enter to proceed.
29 Having decanted all supernatant, move the 48-well plate to position B2.
30 Remove the liquid waste container from the deck and set aside for now.
  A. **CAUTION:** Dispose per local guidelines taking care to note that this supernatant contains both biohazardous material (e.g., plasma) and 2-mercaptoethanol.
31 For the 12-column reservoir at position A2, remove the seal from column 3 and part of column 4 and add 20 mL heated lysis buffer to column 3 as prompted by the system. Place the reservoir back at A2.
32 Press enter to proceed. The robot will now add 300 *µ*L of lysis buffer to each row of the 48-well plate and dispose of used tips.
33 Seal 48-well plate. Vortex passing the plate bottom across the vortex to ensure well-mixed samples across it for at least 30 seconds.
34 Incubate the 48-well plate for 10 minutes at 60°C.
35 During the incubation, prepare the filter plate included in the Norgen kit and place it on top of a 2 mL 96-well collection plate. If this is the first 48-well sample batch, seal the entire filter plate and cut between rows 6 and 7 with a razor. Rows 7-12 will remain sealed until processing the next batch of 48 samples. Set this aside for now.
36 Once there is approximately 1 minute left in incubation, remove the seal from column 5 and part of column 6 from the 12-column reservoir at position A2 and add 20 mL 200-proof EtOH to column 5.
37 Remove seal and place the 48-well plate back at position B2.
38 Press enter to proceed. The robot will now add 300 *µ*L of 200-proof to each row of the 48-well plate and dispose of used tips.
39 As the robot proceeds, change the incubator set point as follows.
  A. Processing a total of 96 samples, following 1st set of 48 samples: Keep the incubator at 60°C.
  B. Processing a total of 96 samples, following 2nd set of 48 samples: Preheat the incubator temperature to 37°C for DNase incubation in next section.
  C. Processing a total of 48 samples, following 1st set of 48 samples: Preheat the incubator temperature to 37°C for DNase incubation in next section.
40 Seal 48-well plate and vortex for 1 minute passing the plate bottom across the vortex.
41 Remove the regular length tip box at position E1, place the lid back, and set it aside. You will use this again at step 51.
42 Place an extended length tip box (blue for Rainin) at position E1 without the lid. It is important to use extended length tips as regular length do not reach the bottom of the 48-well plate.
  A. 1st set of 48 samples: Use a new extended length tip box.
  B. 2nd set of 48 samples: Use the same extended length tip box as in A. Make sure that pipette box is positioned such that the 6 remaining tip rows are closest to you when facing the system.
43 Remove the 2-column reservoir at A1 and dispose of it.
44 Place an empty tip box at position C1. This will act as trash for this step and the system will dispose of used pipette tips here. Ensure when tips are disposed of that they fall flat and do not obstruct robotic degrees of motion.
45 Place the Norgen filter plate you prepared in step 35 at position A1 with the corner well A1 closest to you on the left when facing the system.
  A. **CAUTION:** Often there is a little wiggle room in how the filter plate sits on top of the collection plate. Take care to always position the filter plate in the same spot relative to the collection plate prior to running the protocol so that the pipette’s saved calibration remains true. We recommend pulling the filter plate as far up and to the left when facing the system as the collection plate allows each time.
46 The system will now prompt you and ask what filter row to start at.
  A. 1st set of 48 samples: Type 1 and press enter.
  B. 2nd set of 48 samples: Type 7 and press enter.
47 Vortex the 48-well plate again focusing on the first column. Remove the seal from that column alone using a razor blade as before. Place at position B1.
  A. **CRITICAL:** Move quickly after vortexing so that the slurry mixture does not settle again.
48 Press enter to proceed. The system will now pick up long pipette tips, aspirate the sample and reagents present in column 1 of the 48-well plate in B1, transfer it to column 1 of the 96-well filter plate in A1, and dispose of used tips in C1.
49 Repeat steps 47 and 48 for each row of the 48-well.
50 Centrifuge filter and collection plate with sample rows unsealed (i.e., rows 1-6 for 1st 48-well plate and then rows 7-12 for 2nd) for 4 minutes at 3900 RPM.
51 Place regular length tip box (gray for Rainin) set aside at step 41 at position E1 in the same orientation (i.e., empty rows facing the front). The system knows to start picking up tips at row 6. Set aside extended length filter tip box with lid (blue for Rainin); you will use these extended length tip rows for the next set of 48 samples.
52 Following the system’s prompting, place a 2-column reagent reservoir at position A1. Place the reservoir in A1 such that the open, clean reservoir boat is that closest to the front of the system and user.
  A. 1st set of 48 samples: Use a new 2-column reservoir. Like before, seal both reservoir boats and then remove the seal only from one.
  B. 2nd set of 48 samples: Use the same reservoir as in option A. Make sure to flip it such that the previously sealed, clean boat is now facing the front.
53 Place filter plate with collection plate underneath at position B1.
  A. 1st set of 48 samples: Filter plate column 1 should be closest to the front of the system and user.
  B. 2nd set of 48 samples: Filter plate column 12 should be closest to the front of the system and user.
54 Add 24 mL of Norgen wash buffer to the open reservoir boat in the 2-column reservoir at A1.
55 Press enter to proceed. The system will now add 400 *µL* of wash buffer to the 6 rows closest to the front and dispose of the used tips.
56 Centrifuge the plate with unsealed for 3 minutes at 3900 RPM taking care to ensure the system is balanced. Discard flow through.
57 Repeat steps 53 to 56 twice more for a total of 3 washes.
58 Centrifuge filter and collection plate with sample rows unsealed (i.e., rows 1-6 for 1st 48-well plate and then rows 7-12 for 2nd) for an additional 5 minutes at 3900 RPM to ensure it is fully dry.
59 Replace collection plate underneath filter plate with the elution plate. Take care to match positions between the filter and elution plate such that filter position A1 sits on top of elution plate position A1.
  A. 1st set of 48 samples: Use a new, labeled 96 deep well plate.
  B. 2nd set 48 samples: Take the previously used elution plate out of the fridge, remove the seal, and place underneath the filter plate.
60 For the 12-column reservoir at position A2, remove the seal from column 9 and part of column 10 and add 10 mL elution buffer to column 9 as prompted by the system. Place the reservoir back at A2.
61 Place filter plate with elution plate underneath at position B2.
  A. 1st set of 48 samples: Filter plate column 1 should be closest to the front of the system and user.
  B. 2nd set of 48 samples: Filter plate column 12 should be closest to the front of the system and user.
62 Press enter to proceed. The system will now add 120 *µ*L of elution buffer to the first 6 rows and dispose of the used tips. In our hands, adding 120 *µ*L of elution buffer results in a final elution volume of 100 *µ*L.
63 Centrifuge filter and elution plate with sample rows unsealed (i.e., rows 1-6 for 1st set of 48 samples and then rows 7-12 for 2nd set of 48 samples) for 5 minutes at 3900 RPM.
64 Confirm that 6 sample columns in elution plate now contain 100 *µ*L of flow-through and eluted cfRNA.
  A. Processing a total of 96 samples, following 1st set of 48 samples
    i. Move the filter plate to a new collection plate and set aside keeping columns 7-12 sealed.
    ii. Seal the elution plate that now contains eluted samples for columns 1-6 and place in the fridge.
    iii. Set aside tip box with 3 rows remaining. You will use these in “RNA cleaning and concentration” section.
    iv. Enter 0 at the system’s prompting when asked whether to “`Proceed_to_Zymo?`”, which will end the script. Repeat steps 4-63 for the 2nd set of 48 samples.
  B. Processing a total of 96 samples, following 2nd set of 48 samples
    i. Enter 1 at the system’s prompting when asked whether to “`Proceed_to_Zymo?`”.
    ii. Set aside tip box with 3 rows remaining. You will use these in “RNA cleaning and concentration” section.
    iii. Proceed to next section, “DNase digestion”.
  C. Processing a total of 48 samples, following 1st set of 48 samples
    i. See steps 64(b)i and 64(b)iii under Option 64b

### DNA digestion

Timing: 20 minutes

65 Prepare a master mix of DNase and 10x DNase buffer. Each sample requires 11 *µL* of buffer and 2 *µL* of DNase. For 96 samples, combine 1320 *µL* buffer and 240 *µL* DNase enzyme. Mix well and keep on ice until ready.
66 Dispense 13 *µL* of DNase master mix by hand to each sample well using a P20 multichannel pipette and a disposable reagent reservoir.
67 Seal and mix by vortexing lightly.
68 Incubate for 20 minutes at 37°C. During the incubation, proceed to preparation steps for next section.

### RNA cleaning and concentration

Timing: 1 hour

68 Place a tip box at position E1. If you are processing a total of 96 samples, use the tip box you set aside in step 64 with 3 rows remaining facing the front. If you are processing a total of 48 samples, use a new tip box.
69 Seal and place two new 2-column reagent reservoirs at positions A1 and D1.
70 Remove the seal from the column closest to you when facing the system for the D1 reservoir.
71 Prepare the filter plate included in the Zymo kit and place it on top of a 2 mL 96-well collection plate. Set this aside for now.
72 Once the incubation is complete, place the sample plate at position C1.
73 Enter either 6 or 12 at the system’s prompting to indicate the final row number to be processed where 6 would indicate 48 samples and 12 would indicate 96 samples.
74 As prompted by the system, add 27 mL binding buffer to the unsealed reservoir column in position D1.
75 Press enter to proceed. The system will now add 226 *µ*L of binding buffer to the number of specified rows and dispose of the used tips.
76 As prompted by the system, remove the seal for the second reservoir column in position D1 and add 50 mL 100% ethanol.
77 Press enter to proceed. The system will now add 339 *µ*L of ethanol to the number of specified rows and dispose of used tips.
78 If processing 96 samples, change the tip box for a new tip box as prompted by the system.
79 Place the filter plate prepared at step 71 at position B1.
80 Ensure that corner A1 for both the sample plate at C1 and filter plate at B1 is top left closest to the user when facing the system.
81 Press enter to continue. The system will now sequentially mix each row of samples and then transfer the full sample volume to the corresponding filter plate row switching tips between rows.
82 Once all samples have been successfully transferred to the filter plate, centrifuge the now-filled filter plate for 5 minutes at a speed between 3000-5000xg. Ensure the centrifuge is balanced. Dispose of flowthrough and reassemble the filter and collection plate in the same orientation.
83 Add 48 mL of RNA preparation buffer to the first reservoir boat facing the user in the 2-column reservoir at A1.
84 Press enter to proceed. The system will now add 400 *µ*L of RNA preparation buffer to rows and dispose of the used tips.
85 Centrifuge the plate unsealed for 5 minutes at a speed between 3000-5000xg taking care to ensure the system is balanced. Discard flow through and reassemble filter and collection plate.
86 Add 84 mL of wash buffer to the second reservoir boat facing the user in the 2-column reservoir at A1.
87 Press enter to proceed. The system will now add 700 *µ*L of wash buffer to rows and dispose of the used tips.
88 Centrifuge the plate unsealed for 5 minutes at a speed between 3000-5000xg taking care to ensure the system is balanced. Discard flow through. Using a new collection plate, reassemble filter and collection plate.
89 Add 48 mL of wash buffer to the second reservoir boat facing the user in the 2-column reservoir at A1.
90 Press enter to proceed. The system will now add 400 *µ*L of wash buffer to rows and dispose of the used tips.
91 Centrifuge the plate unsealed for 5 minutes at a speed between 3000-5000xg taking care to ensure the system is balanced. Discard flow through and dispose of collection plate. Place filter plate on a new 96-well PCR plate, the elution plate, taking care to align the wells such that the numbers match.
92 Dispense 12.5 *µ*L of NF-H2O to each sample well by hand using a P20 multichannel pipette and a disposable reagent reservoir taking care to wet the filter only and not splash the sides.
93 Centrifuge the plate unsealed for 5 minutes at a speed between 3000-5000xg taking care to ensure the system is balanced.
94 During the centrifugation, label two 96-well PCR plates, each of which will be used for a separate preparation of the RNA, and 2 8-well PCR strips, both of which will be used to confirm RNA quality on the Bio-analyzer. Place both 96-well PCR plates in cold plate holders.
95 After centrifugation, confirm that cfRNA has been eluted into wells as expected. Place elution plate in a cold plate holder. Pipette 4 *µ*L of eluted cfRNA into the corresponding well of each labeled plate. Seal the labeled plates and place in −80°C freezer. Pipette 1 *µ*L of eluted cfRNA from randomly selected wells across plate (see General setup) to be run on the Bioanalyzer. Cap PCR-strips and seal original elution plate. Freeze all at −80°C.

### Downstream processing

#### RNA quality check

Timing: 1 hour

96 Prepare samples and RNA Pico chip for Bioanalyzer per manufacturer instructions. In our experience, we have found it easier to pipette in 5 *µ*L of RNA 6000 Pico marker into the 8-well PCR strip containing 1 *µ*L of sample, mix, and then transfer a total of 6 *µ*L into the corresponding chip well as compared to pipetting marker and sample sequentially. Avoid introducing bubbles in this process.
97 Assess sample quality following run. Typically, we expect anywhere between 0.1-10 ng of total RNA extracted from 1 mL of plasma.

#### RT-qPCR following Option A (Exogenous RNA control)

Timing: 2 hours

98 Prepare master mix for iTaq Universal Probes One-Step Kit per manufacturer instructions.
99 Combine 1 *µ*L of isolated exogenous RNA with 0.5 *µ*L of the corresponding Taqman probes and 8.5 *µ*L of master mix.
100 Perform real time RT-qPCR on the Biorad CFX system (either 96 or 384) using the following program: 50°C for 10 minutes, 95°C for 1 minute, 95°C for 10 seconds, 60°C for 20 seconds, Return to 95°C for 10 seconds followed by 60°C for 20 seconds for a total of 40 cycles.
101 Obtain cycle thresholds from the corresponding CFX software and compare with expected results.

#### RNAseq library preparation following Option B (Frozen plasma samples)

Timing: 2 days

102 Prepare cfRNA sequencing libraries following manufacturer instructions for SMARTer Stranded Total RNAseq Kit v2 using 4 *µ*L of eluted cfRNA.
103 Barcode samples using unique dual index barcodes.
104 Pool in an equimolar fashion and sequence in a paired-end fashion (2×75 bp) to an average depth of 50 million reads per sample.

### Data analysis

106 Clone bioinformatics github repository - https://github.com/miramou/cfRNA_pipeline using the command `git_clone`. Note that this pipeline relies on High Performance Computing (HPC) cluster and large file systems for performance. Do not run locally.
107 Follow instructions in `README` file to get started and set up new project. Note that repo may need to be tweaked given your HPC and job manager. You will ultimately have project specific data saved in two places - one folder (i.e., the run folder) will be home to files that contain run parameters and another will contain raw data and output files from running the pipeline.
108 Within the `root_dir`, create a project specific output folder e.g. `root_dir/project_id/`
109 Download raw reads from the cloud (e.g., AWS, GCloud, etc.) into a sub-directory e.g. `root_dir/project_id/raw_data/`
110 Create an analysis specific directory e.g. `analysis/`. This is where all the pipeline specific files will be saved like the output from STAR, htseq, etc.
111 Create a `sample_metadata` file. See github for more details and an example of what this file entails - https://github.com/miramou/cfRNA_pipeline/tree/master/run/example.
112 Create a run specific folder that contains `config.yaml` and `run_snakemake.sh` to launch the pipeline and specify project specific parameters like whether reads are single or paired end.
113 Launch a dry run of your data analysis using the bash command:
  · `bash_run/project_name/run_snakemake.sh` where `project_name` indicates a user specific name for their project. We recommend specifying a sample metadata file that contains a subset of all samples if this is your first time to confirm the pipeline runs all the way through.
114 Confirm the number of jobs launched looks right relative to the number of samples in the metadata sheet.
115 Launch the pipeline locally using the argument `snakemake` or launch a master job (better if you have more samples) using the argument `sbatch`.
116 Analyze counts tables as desired. An example analysis is provided in https://github.com/miramou/pe_cfrna.

### Troubleshooting

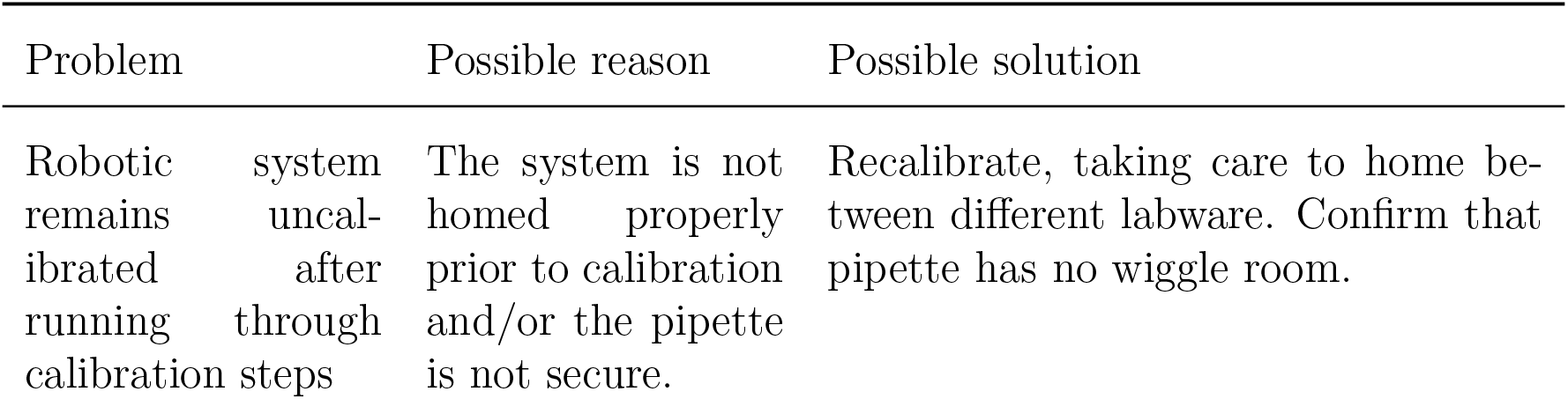
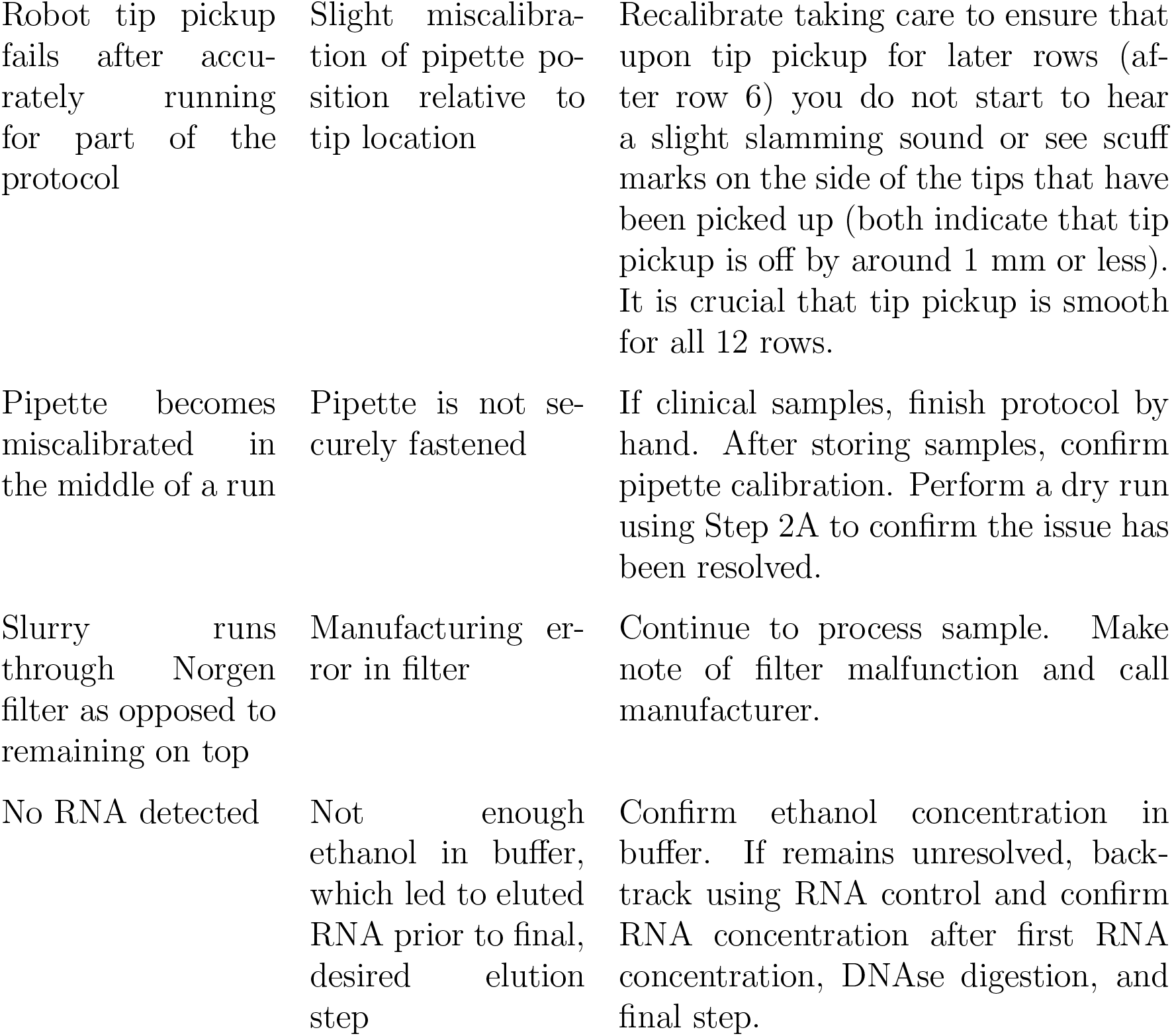
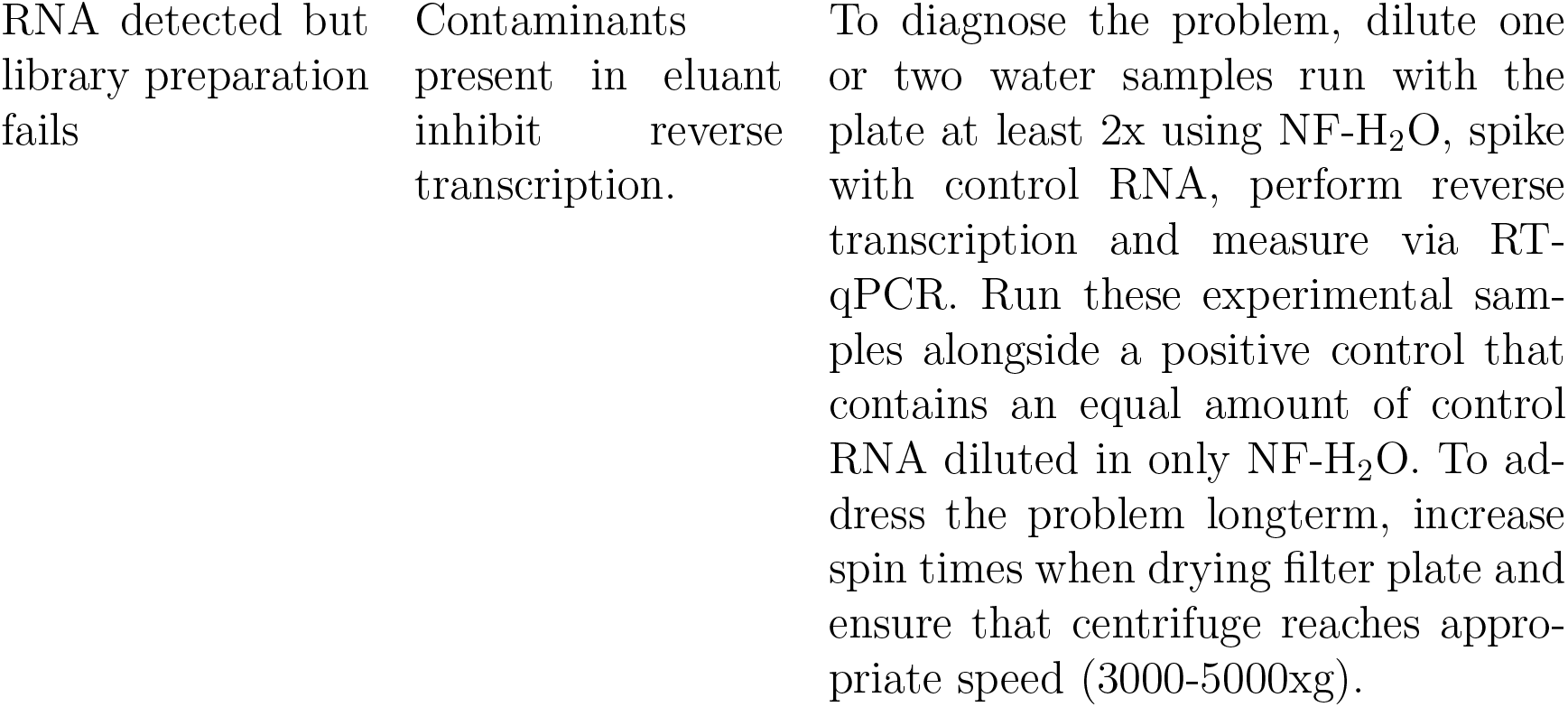

| Problem | Possible reason | Possible solution |
| --- | --- | --- |
| Robotic system remains uncalibrated after running through calibration steps | The system is not homed properly prior to calibration and/or the pipette is not secure. | Recalibrate, taking care to home between different labware. Confirm that pipette has no wiggle room. |
| Robot tip pickup fails after accurately running for part of the protocol | Slight miscalibration of pipette position relative to tip location | Recalibrate taking care to ensure that upon tip pickup for later rows (after row 6) you do not start to hear a slight slamming sound or see scuff marks on the side of the tips that have been picked up (both indicate that tip pickup is off by around 1 mm or less). It is crucial that tip pickup is smooth for all 12 rows. |
| Pipette becomes miscalibrated in the middle of a run | Pipette is not securely fastened | If clinical samples, finish protocol by hand. After storing samples, confirm pipette calibration. Perform a dry run using Step 2A to confirm the issue has been resolved. |
| Slurry runs through Norgen filter as opposed to remaining on top | Manufacturing error in filter | Continue to process sample. Make note of filter malfunction and call manufacturer. |
| No RNA detected | Not enough ethanol in buffer, which led to eluted RNA prior to final, desired elution step | Confirm ethanol concentration in buffer. If remains unresolved, back-track using RNA control and confirm RNA concentration after first RNA concentration, DNase digestion, and final step. |
| RNA detected but library preparation fails | Contaminants present in eluant inhibit reverse transcription. | To diagnose the problem, dilute one or two water samples run with the plate at least 2x using NF-H<sub>2</sub>O, spike with control RNA, perform reverse transcription and measure via RT-qPCR. Run these experimental samples alongside a positive control that contains an equal amount of control RNA diluted in only NF-H<sub>2</sub>O. To address the problem longterm, increase spin times when drying filter plate and ensure that centrifuge reaches appropriate speed (3000-5000xg). |

## Timing

Steps 1-2, Prepare samples: 10 minutes

Steps 3-63, RNA extraction: 3 hours (1.5 hours per 48-well plate)

Steps 64-67, DNA digestion: 20 minutes

Steps 68-95, RNA cleaning and concentration: 1 hour

Steps 96-97, RNA quality check: 2 hours

Steps 98-104, Downstream processing: 1-3 days

Steps 105-116, Data analysis: 1-3 days to yield counts table

## Anticipated results

This semi-automated protocol allows the user to extract cfRNA from 96 samples and confirm sample quality in under 1 day. We previously used this protocol to extract cfRNA from 404 samples in our work [1] and detail sample quality metrics in Extended Data Figure 1. Extracting cfRNA and confirming sample quality from all samples required 27 hours (1 96 well batch was processed over 2 days because of when samples were received). Preparing libraries for sequencing took another 10 days. Altogether, we prepared sequencing libraries for all 404 samples within 2.5 weeks as compared to the months that it would have required to process samples manually. We then processed this raw data as detailed previously (see https://github.com/miramou/pe_cfrna), performing differential expression, building a machine learning model, and finally understanding the tissue and cellular contributors to cfRNA.

Overall, we have shown that semi-automated sample processing can be performed affordably and quickly within individual academic labs. Shifting to semi-automated sample processing alleviates a significant bottleneck in discovery-focused work and allows for seamless transitions from sample collection to data generation and analysis. With this robust and scalable solution, we can test hypotheses faster and spend less time on repetitive but important tasks like sample processing.

## Supporting information

Supplementary Movie, Also available on youtube

## Data availability

Source data can be found in the zip file - `data.zip`.

## Code availability

All software, including the data analysis required to generate all figures, are available at https://github.com/miramou/cfRNA_pipeline_automation and https://github.com/miramou/cfRNA_pipeline.

## Acknowledgements

We are deeply grateful to both Norma Neff and Rene Sit for their sequencing expertise. We also thank Brian Yu for helping brainstorm and providing indispensable advice around how to best use the Opentrons system. Figures 1A and 3A were created with `BioRender.com`. This work was supported by the Chan Zuckerberg Biohub. M.N.M. is supported by the Stanford Bio-X Bowes Fellowship.

## Author contributions

M.N.M. conceptualized and designed this protocol and collected and analyzed the data in collaboration with S.R.Q.. All authors contributed to the writing and editing the manuscript.

